# Long-term evolution of prokaryotic genomes in a chemolithotrophic cave over 5.5 million years of isolation

**DOI:** 10.1101/2025.04.10.648229

**Authors:** Zaki Saati-Santamaría, Alena Nováková, Lorena Carro, Fernando González-Candelas, Jaime Iranzo, Alexandra Maria Hillebrand-Voiculescu, Daniel Morais, Tomáš Větrovský, Petr Baldrian, Miroslav Kolarik, Paula García-Fraile

## Abstract

Fluctuating conditions drive adaptive evolution, yet understanding how genomes evolve under stable conditions over extended periods remains a major challenge, since most research on microbial evolution relies on short-term experiments or phylogenetic comparisons. The Movile Cave, isolated from external influences 5.5 million years, offers a unique opportunity to explore microbial evolution under prolonged environmental stability. Here, we analyzed metagenome-assembled genomes from this cave, revealing that prokaryotes exhibit lower gene diversity and higher levels of pseudogenization compared to those from non-isolated environments, mainly affecting housekeeping functions involved in translation. Functional redundancy across genomes remained comparable to related habitats. Our results suggest that pseudogenization may serve as a fine-tuning mechanism to reduce excess redundancy. Although horizontal gene transfer is limited overall, the cave virome seems to contribute to microbial adaptation through the transfer of auxiliary metabolic genes. Movile microorganisms harbor fewer phage-defense systems than counterparts in related environments, suggesting a long-term adaptation to a relatively stable virosphere. Our findings indicate that prolonged isolation under stable selective pressures does not necessarily lead to major genomic divergence, but rather promotes adaptive gene loss. This study provides key insights into how long-term stability shapes microbial genome evolution and ecosystem function.

## Main

Microorganisms are known to inhabit nearly all Earth environments, where they play essential biogeochemical roles, such as recycling nutrients (e.g., C, N, P, S) and modulating the performance of hosts (animals and plants) [1,2]. To ensure their survival, they must undergo adaptive evolution that make them fitter for the habitats where they dwell. These include nucleotide substitutions, gene duplications and losses, and even horizontal gene transfer (HGT) events involving large DNA regions. The latter has been suggested as the primary driving force in the evolution of certain microorganisms [3,4]. However, the evolutionary forces driving gene loss and genome streamlining remain debated. In some cases, genome reduction appears to be an adaptive response to stable or nutrient-limited environments, where selection favors a more efficient genetic repertoire. In other cases, it may be largely non-adaptive, resulting from relaxed selection, a general deletion bias, or shifts in mutation rates [5–8]. While gene loss can be facilitated by relaxed selection, it may also reflect an adaptive process when the elimination of certain functions provides a fitness advantage [9,10]. Pseudogenization often results from weakened selective constraints, and may also represent an adaptive transition when gene inactivation confers a selective advantage under specific environmental conditions. In support of this, multiple studies suggest that pseudogenization and genome streamlining [11] can be an adaptive strategy, particularly in contexts where certain metabolic functions become unnecessary [12–17]. Understanding the mechanisms and evolutionary pressures shaping adaptive gene loss is therefore crucial to deciphering microbial adaptation to the environment.

Numerous comparative genomic studies have been conducted among microbial taxa from different habitats, with the aim of describing genomic divergences associated with niche occupancy or phenotypic traits [18–20]. Additionally, studies employing laboratory experiments to track microbial adaptive evolution under external pressures further emphasized the utility of incorporating time (prospective) as a variable [21,22]. The long-term evolution experiment with *Escherichia coli*, started on 1988, illustrates perfectly the complexity of prokaryotic evolution, and the relevance of perspectives that consider long timeframes [23–25]. Also, the recovery of preserved DNA from frozen samples can be used to investigate ancient genomes, which help to establish reference points for life evolution [26]. Although the study of microbial evolution has made significant strides, there are still gaps in exploring evolutionary dynamics in specialized environments [1,27], such as those with long-term ecological stability, where innovative approaches can shed new light on adaptive processes.

Genome evolution events are typically driven by the need to adapt to ever changing selective pressures, such as transitioning to a different ecological niche with distinct nutrient resources, combating the effects of antibiotics, or increasing the competitive fitness [4,21]. However, questions arise regarding the long-term evolution of life in the absence of such shifts on the effects of selective pressures. For instance, to what extent do microorganisms retain new genes acquired through horizontal gene transfer (HGT) events when these genes are no longer necessary to cope with novel stresses? To what degree do microorganisms undergo pseudogenization of once-useful genes, and how does this contribute to genome streamlining and adaptation to these more stable conditions?

The Movile Cave in Romania represents a unique chemolithotrophic-based environment that offers an opportunity to address these questions and study genome evolution events under prolonged, constant environmental conditions. This cave has been sealed from the external environment, including outer life, and climatic shifts, for ∼5.5 million years (Myr) [28]. Consequently, life within the cave has not undergone significant shifts in the presence of selective pressures. It is plausible that the evolutionary trajectory of life inside the cave differs from that outside. Since its discovery in 1986, the cave has yielded dozens of endemic novel species, including invertebrates, bacteria, and fungi [28–30]. Due to the high levels of H_2_S, CH_4_ and NH_4_ ^+^ in the underground water, along with elevated H_2_SO_4_ and CO_2_ but low O_2_ levels in the atmosphere, and complete absence of sunlight, this ecosystem thrives on the energy acquisition and nutrient recycling activities of metanotrophs, methylotrophs, and sulfur-oxidizing microorganisms, resembling the microbial communities found in deep-sea hydrothermal vents. Consequently, researchers have primarily focused on studying the chemolithotrophic characteristics of microbial life within this cave [28,31–35].

Here, we used the prokaryotic DNA from several locations of the Movile Cave with the aim to investigate microbial genome evolution. We hypothesize that the rate of evolution is slower in the cave due to (I) the limited number of interactions with invading species,, which reduces the influx of novel genetic material, and (II) the stable selection pressure resulting from the isolation of the cave. Finally, as the virome of the Movile Cave has been largely neglected, we analyzed its composition, and their likely ecological roles through the investigation of their auxiliary metabolic genes (AMGs). By investigating the prokaryotic genomes, our study aims to provide a deeper understanding of how long-term isolation, stable selective pressures, and viral interactions shape microbial evolution in isolated ecosystems.

## Results

### Recovering high-quality metagenome-assembled genomes from putative novel taxa

The Movile Cave ecosystem has remained isolated from external influences for approximately 5.5 Myr. Hence, we aimed to analyze the prokaryotic genomes found in this environment to gain insights into the evolutionary processes that these microbes have undergone in this lithotrophic ecosystem, where significant environmental changes, acting as selective pressures, have been absent. We obtained a total of 111 bins (>50% completeness, <10% contamination) from diverse binning algorithms (Supplementary Table 1). These bins were taxonomically classified into 12 bacterial phyla, and one archaeal phylum (*Thermoplasmatota*) (Supplementary Table 1). Out of these bins, we obtained 21 metagenome-assembled genomes (MAGs) (>90% completeness, <5% contamination). Of these, 6 resulted from the de-replication and quality improvement process using the DASTool algorithm (Supplementary Figure 1, Supplementary Table 1).

We compared the MAGs with reference genomes from validated type strains and found that only one could be classified within a known species (*Sulfurospirillum deleyianum;* dDDH: 70.90%; ANIb: 96.5%; dastool_metabat2.18). According to the phylogenomic analyses and genome similarities, four of the remaining MAGs (dastool_concoct_19, dastool_maxbin2_145, dastool_metabat2_12, and dastool_metabat2.39) are undetermined species of known genera (*Edaphobacter, Desulfuromonas*, *Rummeliibacillus*, and *Sedimentibacter,* respectively), and one of them (dastool_metabat2.89) seems to be a member of a undetermined genus in the phylum *Actinomycetota* (Supplementary Data). Yet, due to the lack of reference genomes for all these taxa, we just propose the description of the novel species “*Candidatus* Edaphobacter cavernae” and “*Candidatus* Sedimentibacter cavernae” (Supplementary Data; ‘Protologue’). Finally, the MAG named as ‘dastool_metabat2_14’ presents very low similarities with members of different phyla (4% dDDH, approximately), which indicates that this taxon is largely different from any yet described bacteria and makes classification impossible even at upper taxonomic levels (Supplementary Data).

All the above-mentioned MAGs were detected in airbell II, whose atmosphere is depleted in O_2_ (7-10%) and has high CO_2_ (2-3.5%) and CH_4_ (0.5-1%) levels [35,36], but the genome of the novel species “*Candidatus* Edaphobacter cavernae” was binned from airbell I (6.18% abundance), which, unlike airbell II, has not so restrictive conditions (1.5% CO_2_, 19% O_2_) [36] (Supplementary Table 2).

### Gene loss drives adaptation of Movile cave bacterial genomes

Our comparative analysis with the genomes and MAGs of external strains (Supplementary Table 3) revealed that MAGs from the cave possessed fewer unique genes (singletons) compared to their counterparts from different environments (Figure 2a, Supplementary Figure 2, Supplementary Figure 3), except in the case of the *Edaphobacter* pangenome (Supplementary Figure 4). This suggests that the cave’s microorganisms, likely subjected to lesser shifts on selective pressures, have retained fewer accessory genes. Notably, the unique genes within the cave’s genomes were primarily associated with ’Cell wall/membrane/envelope biogenesis’, ’Inorganic ion transport and metabolism’, and ’Defense mechanisms’, among others (Figure 2a, Supplementary Figure 2, Supplementary Figure 3, Supplementary Figure 4). Genes related to inorganic ion transport and metabolism are particularly relevant in the context of a chemolithotrophic cave environment, where microorganisms likely rely on the oxidation of inorganic compounds as primary energy source. The enrichment in Cell wall/membrane/envelope biogenesis genes may reflect adaptations for enhanced cell envelope stability under extreme cave conditions, such as low nutrient availability or exposure to toxic metals, helping to retain essential ions and prevent leakage. Similarly, genes related to Defense mechanisms suggest that, in a nutrient-limited ecosystem, microbes may engage in competition by producing antibiotics or inhibitory compounds. Together, the functional categories highlight the potential adaptation of cave bacteria to their unique ecological niche, suggesting that these microorganisms prioritize cellular integrity, defense, and the utilization of inorganic compounds for survival in that chemoautotroph-driven ecosystem.

**Figure 1.**
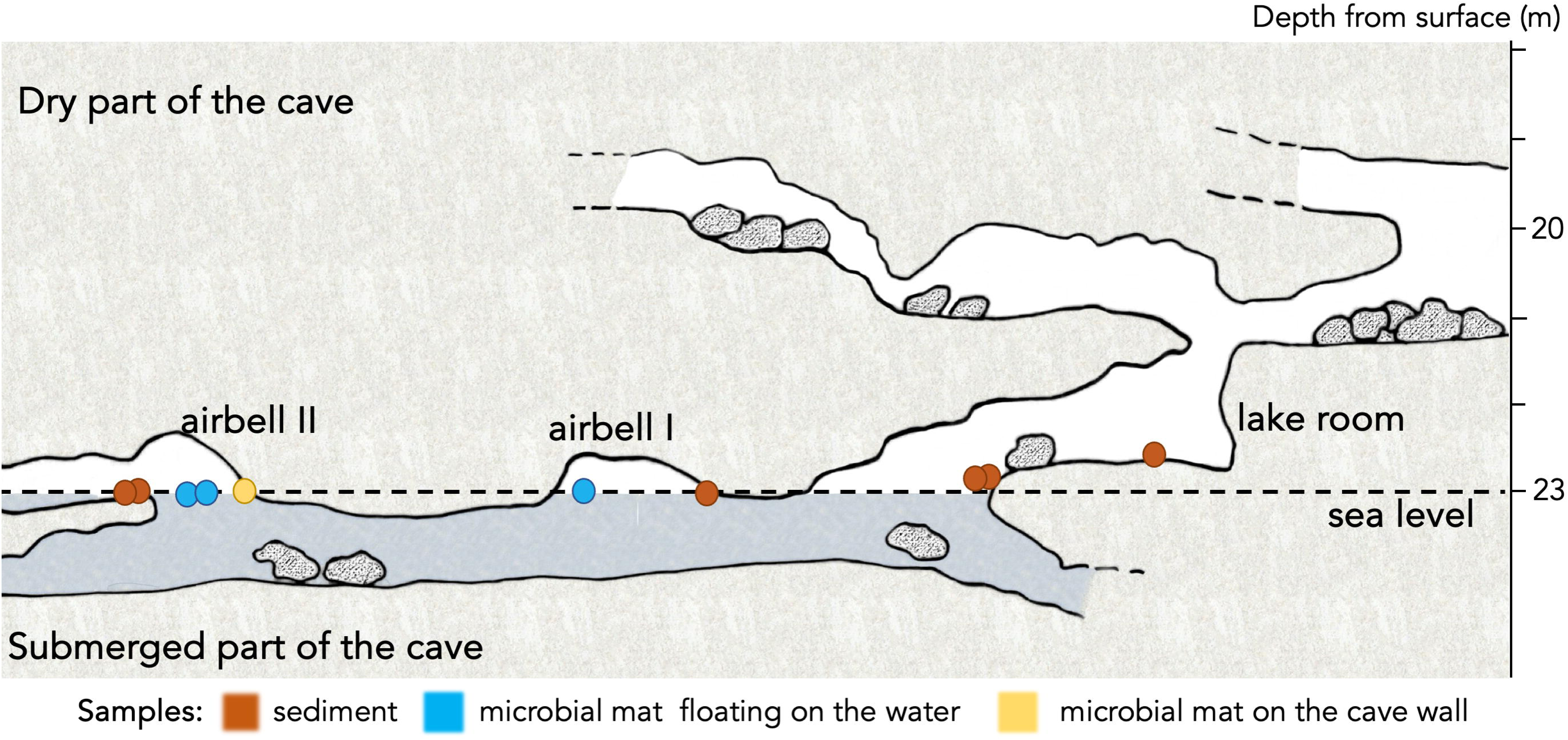
Map of the sampling sites in Movile Cave (Romania). The circles represent samples used for shotgun metagenomic analyses and are colored according to the sample type. This map has been redrawn from the original available at http://old.lefo.ro/iwlearn/movile/maps.htm.

**Figure 2.**
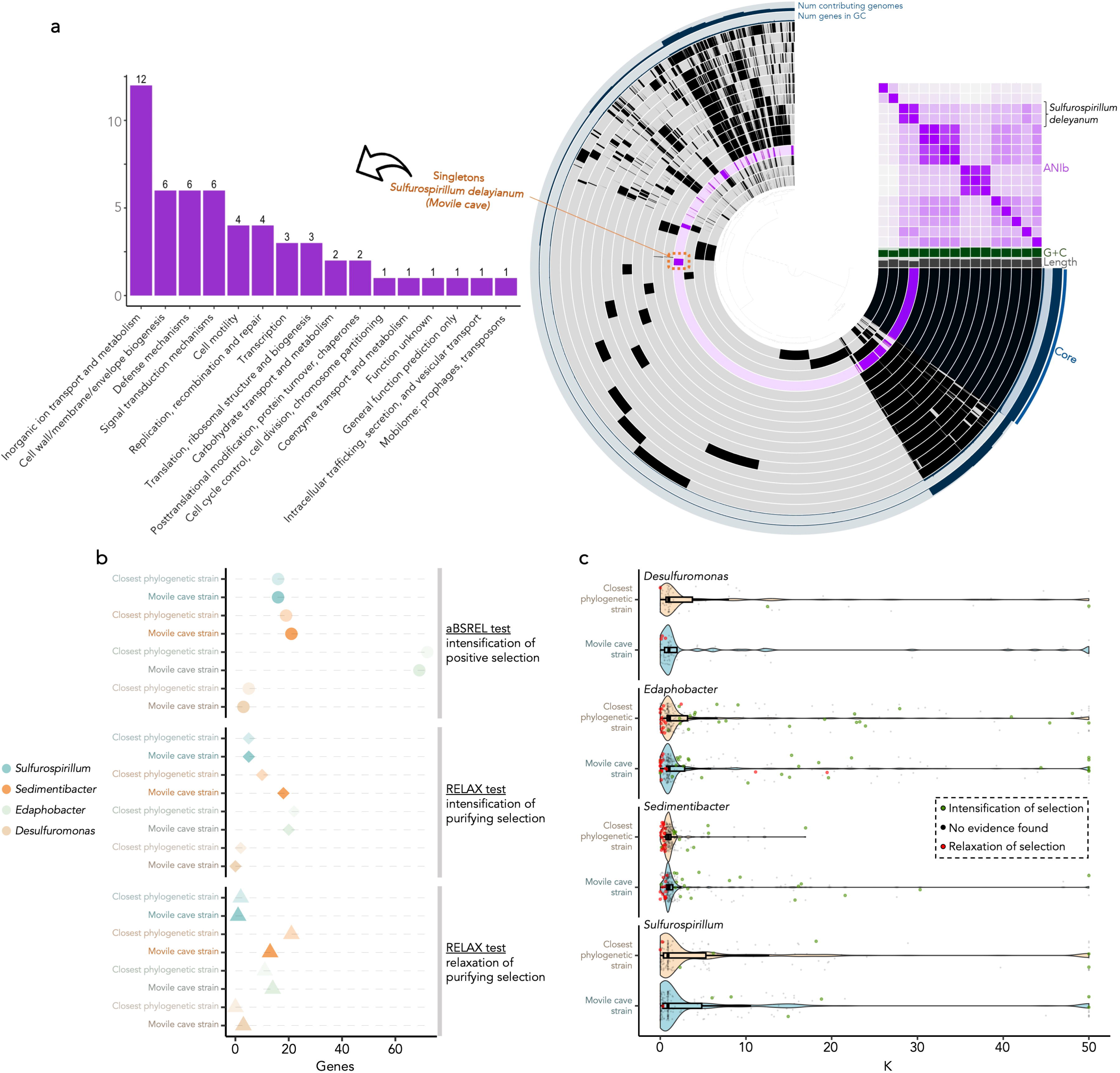
Comparative genomic analysis and selection dynamics of prokaryotic MAGs from Movile Cave and related environments. a) Comparative genomic analysis of the *Sulfurospirillum deleyianum* metagenome-assembled genome (MAG) from Movile Cave (*dastool_metabat2.18*) alongside genomes and MAGs from other environments within the same genus (Supplementary Table 3). Each circle represents the presence (dark color) or absence (light color) of gene clusters within each genome. The purple circle indicates the data from the cave MAG. Shared ANIb values across all genomes are depicted in the purple heatmap. On the left side, the functional categories of the singletons found in the *Sulfurospirillum deleyianum* MAG from this study are shown. This summary highlights the metabolic traits that likely drove the limited ecological speciation of this species within this environment. Notably, this genome exhibits the lowest count of singletons among all genomes within this genus. b) Results of the HyPhy selection analyses on Movile Cave prokaryotic MAGs. The analysis was conducted using all available public genomes and bins from the same genera as references. Core genes (80% nucleotide identity and coverage thresholds) were aligned and then analyzed with RELAX and aBSREL tests. The figure shows the number of core genes under intensification or relaxation of selection (p < 0.05) in the branches of each tested Movile MAG. For comparison, the same analyses were performed on the most phylogenetically close genome. c) Intensity of selection on core genes (K parameter; RELAX test). Genes with significant intensification or relaxation of selection (p < 0.05) are represented with larger colored dots.

We retrieved the core genes from each of the four pangenomes mentioned above (*Edaphobacter*, *Desulfuromonas*, *Sulfurospirillum*, and *Sedimentibacter*) to further investigate the evolution of cave-specific genes in comparison to external references. Comparative analyses of genome-wide selection pressures across cave-derived genomes and their phylogenetically closest counterparts revealed similar levels of selection (Figure 2b). This indicates that whether the Movile genome or the most phylogenetically close genome is used as the test or as the reference in the selection analyses, a similar number of genes is identified under selection. This suggests that the cave genomes have been subjected to similar overall levels of selection to those experienced by their closest relatives. However, given the long isolation of the cave, it is unclear whether these selection signals reflect historical adaptation events or more recent ongoing evolutionary processes. Considering the potential lack of fluctuating stressors in the cave acting as selective pressures, our initial hypothesis was that purifying selection could have had a pronounced effect within the cave. However, we found that the pattern of relaxation and intensification of this purifying selection is similar in both sets of genomes (Figure 2c).

To further assess the functional consequences of selective pressures in the cave genomes, we also compared the ratio of pseudogenes in our bins to that on bins from other related locations (Supplementary Table 4). We found a noteworthy increase of pseudogenization in the Movile Cave genomes compared to the others (Wilcoxon rank-sum test, *p* = 2.1×10^-2^) (Figure 3a, Supplementary Figure 5). We also focused on intergenic pseudogenes, as these are more prone to represent decaying gene sequences than other pseudogene categories, which reinforced the evidence that Movile genomes have undergone a higher pseudogenization (Wilcoxon rank-sum test, *p* = 8.5×10^-16^) (Figure 3b).

**Figure 3.**
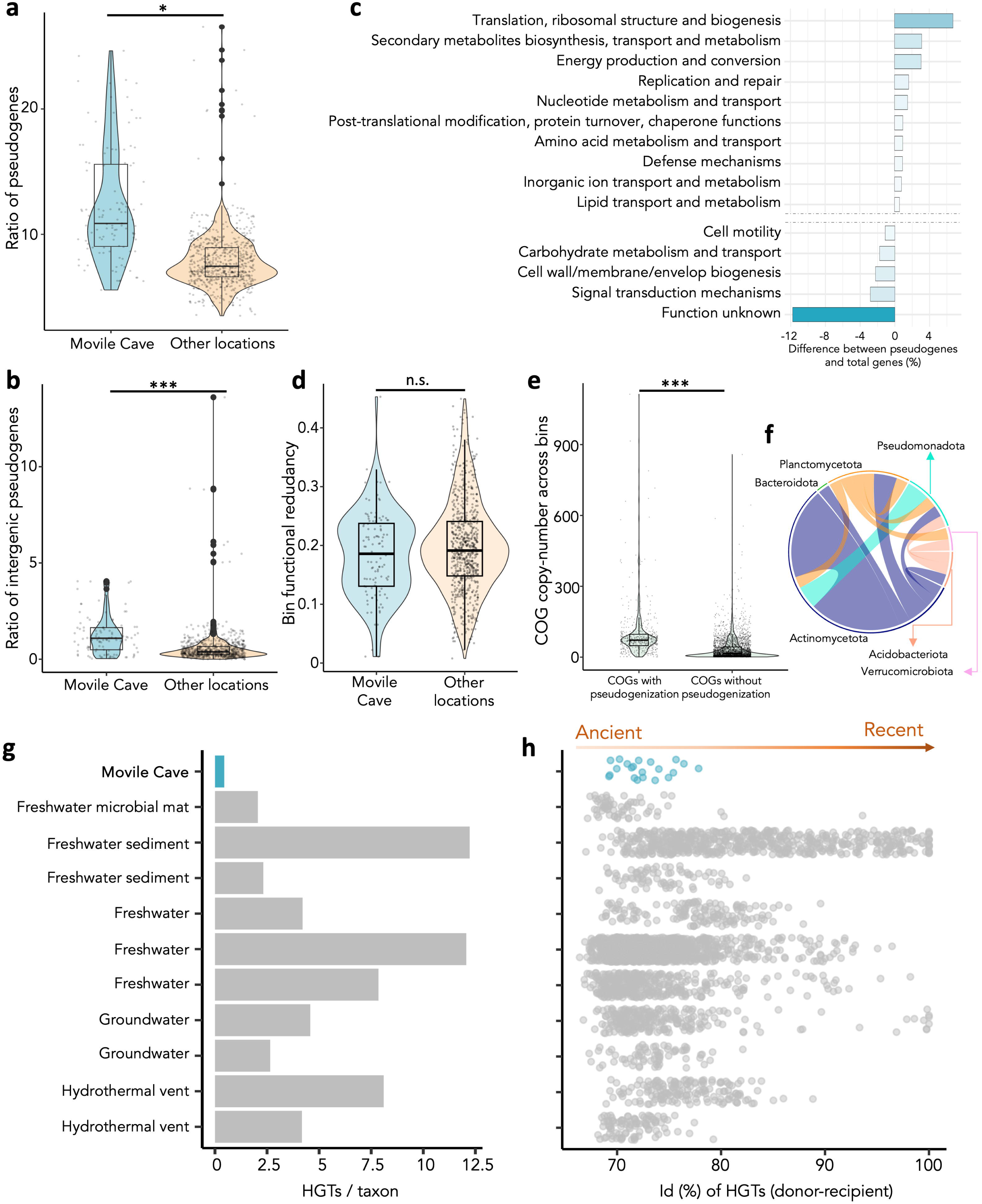
Comparison of evolutionary features between binned genomes from Movile Cave and other environments with similar ecological conditions. a) Ratio of total pseudogenes and b) intergenic pseudogenes, normalized by the total number of CDSs per genome, in bins derived from Movile Cave metagenomes and those from related habitats (Supplementary Table 4). c) Metabolic categories enriched in pseudogenes in the Movile Cave bins. Each bar represents the difference between the proportion of pseudogenes within a specific COG category and the total functions of that category across the entire proteome of the bin. Positive values indicate categories more prone to pseudogenization in the cave. d) Comparison of functional redundancy between cave bins and non-cave genomes. Boxplots show redundancy values for functional categories within the Movile Cave bins and other environmental genomes. No significant differences were observed between the two groups (n.s. = non-significant), suggesting similar levels of functional redundancy despite the cave’s long-term isolation. e) Number of copies of ‘intact’ COG functions in the Movile Cave prokaryotic bins. The boxplots compare the copy number (i.e., redundancy) of COGs that include at least one pseudogenized gene versus those that do not include any predicted pseudogenes. Since pseudogenes cannot be directly annotated, we used the COG annotation of the closest Swiss-Prot homologs employed as references in the Pseudofinder annotation. f) Putative horizontal gene transfer (HGT) events among bins derived from Movile Cave metagenomic data. The circular representation connects pairs of taxa (categorized by phyla) likely sharing horizontally transferred genes (Supplementary Table 3, Supplementary Table 6). g) Proportion of HGTs among bins from Movile Cave and those from other related environments (Supplementary Table 3, Supplementary Table 6). The total number of HGTs was normalized by the total number of input taxa for each location. Raw counts are shown in Supplementary Figure 6. h) Nucleotide similarity values between pairs of genes likely undergoing horizontal transfer. Genes sharing higher nucleotide similarity may have been acquired more recently, whereas those with lower similarity might have evolved over a longer period. Further details on these genes are provided in Supplementary Table 5. \**p* < 0.05; \*\*\**p* < 0.001.

This finding suggests that the observed increase in pseudogenization could reflect adaptive gene loss, where non-essential genes were selectively discarded due to the cave’s unique and isolated environment, leading to genome streamlining. We analyzed the encoded functions of the closest reference intact genes for the intergenic pseudogenes detected in the cave and normalized the number of pseudogenes in each COG category to the total count of intact genes in those categories. Our results indicate that pseudogenization in the cave predominantly affects core functions such as those involved in “Translation, ribosomal structure and biogenesis”, while it has a lesser impact on metabolic and structural processes such as carbohydrate and amino acid metabolism, signal transduction, and cell wall/membrane biogenesis (Figure 3c). Among the top 100 most frequently pseudogenized clusters of orthologous groups (COGs), we observed a notable enrichment of universal bacterial core genes [37], including *rpoB*, *rpoC*, *gyrA*, *recA*, *dnaX*, *secA*, and *tufA*, which are typically conserved and essential for cellular viability (Supplementary Table 5). For instance, *rpoB* (COG0085) and *rpoC* (COG0086), encoding the β and β’ subunits of the RNA polymerase, were pseudogenized in 31 and 32 instances respectively across cave-associated bacterial genomes. Similarly, elongation factors such as *fusA* and aminoacyl-tRNA synthetases like *ileS*, *alaS*, and *leuS* were recurrently disrupted (Supplementary Table 5). These findings indicate an unexpected degree of genomic erosion in functions traditionally regarded as indispensable.

Given the hypoxic conditions of the cave, we specifically examined whether pseudogenization disproportionately affected genes related to oxidative metabolism. Indeed, we found that several key enzymes of the tricarboxylic acid (TCA) cycle and aerobic respiration were pseudogenized, including malate dehydrogenase (*mdh*), fumarate hydratase (*fumC*, *FH*), succinate dehydrogenase (*sdhA*), and 2-oxoglutarate dehydrogenase (*kgd*). Additionally, components of the electron transport chain, such as subunits of NADH dehydrogenase (*nuoD*, *nuoH*, *nuoM*, *nuoG*,) and cytochrome c oxidase (*coxA*, *cydA*), are also pseudogenized (Supplementary Table 5). This pattern suggests a shift away from aerobic respiration as a primary energy-generating pathway, likely favoring anaerobic metabolism or alternative electron acceptors in response to oxygen limitations. Over evolutionary time, such selective gene loss may have contributed to a more specialized metabolic repertoire, consistent with the cave’s environmental constraints.

### Functional redundancy at the community level despite adaptive gene loss

To further evaluate the functional consequences of selective pressures in cave genomes, we also assessed the redundancy of functional categories within and across bins. Functional redundancy, the presence of multiple copies of genes encoding the same function, can buffer microbial communities against environmental fluctuations by ensuring the persistence of key metabolic processes. Given the long-term stability of the Movile Cave environment, we hypothesized that gene redundancy might be reduced in its microbial genomes due to adaptive gene loss. However, our analysis of COG redundancy within individual bins revealed no significant differences between cave genomes and those from other habitats (Wilcoxon rank-sum test; *p* = 0.2142) (Figure 3d), suggesting that functional redundancy at the single-genome level has been maintained despite long-term isolation. When examining inter-bin redundancy—comparing functional overlap across genomes within each metagenome—we found that the Movile Cave microbiome exhibited a high level of functional redundancy (0.94), comparable to values observed in other environments (range: 0.74–0.95).

When comparing the redundancy of COGs that have undergone pseudogenization with those that have remained intact across all bins, we observed significantly higher redundancy in the former group (Wilcoxon rank-sum test; *p* = 1.0×10^-126^) (Figure 3e). This suggests that functions supported by multiple copies are more likely to tolerate gene degradation, while essential functions with low redundancy tend to remain conserved. This trend is maintained across all the different COG categories (Supplementary Figure 6)

### Limited horizontal gene transfer events among Movile cave bacteria

Considering the important role of HGT in evolution, we aimed to identify genes within the assembled MAGs that might have been acquired through HGT events. Given the restricted pool of microbial genes within the cave ecosystem, such events should have occurred either before the cave was sealed or, if later, they would have likely involved interactions among a limited interacting set of microorganisms residing within the cave. Our analysis yielded only 21 putative HGT events among the 111 bins analyzed (Supplementary Table 6). These genes have been likely transferred from one bin (draft microbial genome) within the cave’s ecosystem to another, even spanning different phyla (Figure 3e). To compare the ratio of HGT events in Movile Cave with that of communities from non-isolated niches, we identified such gene transfers among diverse public prokaryotic bins (see Methods) derived from environments such as hydrothermal vents, microbial mats, freshwater, sediments and groundwater. We found that the species from Movile Cave had a lower proportion of HGT events (Figure 3f, Supplementary Figure 7, Supplementary Table 7). Moreover, we have observed a low similarity between homologous gene pairs (donor - recipient) (<80% nucleotide similarity), which suggest that most of these HGT events took place a long time ago, with subsequent gene diversification (Figure 3g). The encoded functions of these few HGT genes are mainly ATPases and proteins involved in nucleotide turnover (Supplementary Table 6). Our findings suggest that horizontal gene transfer has played a limited role in shaping the current genomic landscape of the cave’s microbial community.

### Presence of auxiliary metabolic genes in the cave’s bacteriophages

Bacteriophages can play a pivotal role in modulating prokaryotic evolution and adaptation by both expressing genes with ecological functions, commonly referred to as AMGs, and facilitating the transduction of ARGs. We found that the abundance of all viruses ranged from 0.008% to 0.25% of the metagenomic reads, with considerable dissimilarity observed among samples (Figure 4a). The most abundant viral order across all the cave locations is *Caudovirales*, which typically infects prokaryotes, followed by *Iridoviridae* viruses, known to infect animals, and the giant *Acanthamoeba polyphaga* Mimivirus, which infects amoebas (Figure 4a).

**Figure 4.**
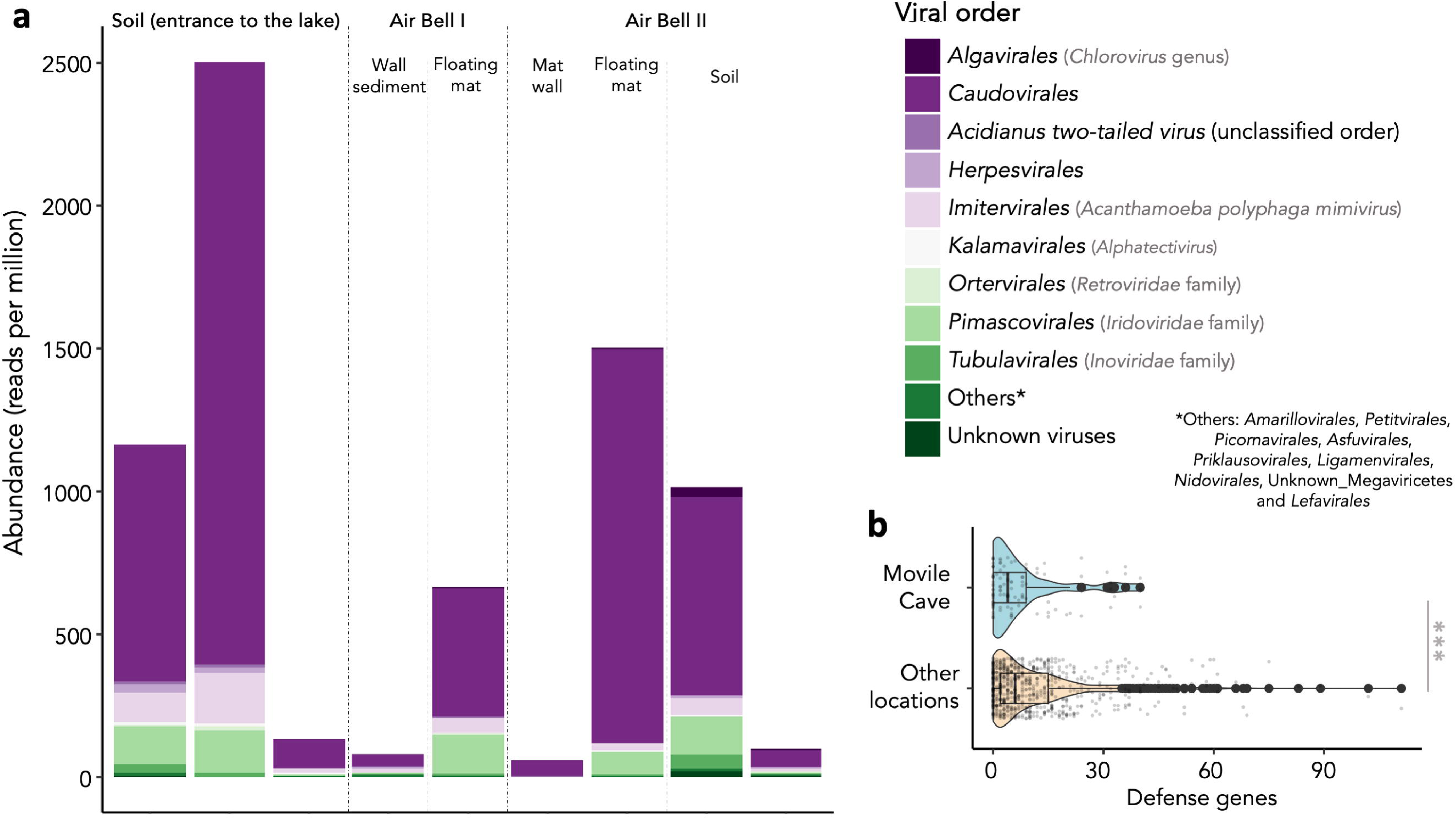
Viral diversity and phage-defense systems in the Movile Cave metagenomes. a) Taxa bar plot displaying the relative abundance of viruses identified in the Movile Cave metagenomes. The abundance of each viral taxon represents the cumulative relative abundance of all contigs classified within that taxon. b) Total count of phage-defense systems identified in the Movile Cave bins and those bins recovered from metagenomes in ecologically related environments. \*\*\**p* < 0.001.

Further investigation into the functions of the encoded proteins within the viral contigs led us to identify some AMGs that could significantly impact microbial ecology. For instance, we detected the presence of a GcrA cell cycle regulator in a viral sequence (o_*Caudovirales*; f_*Myoviridae*; g_*Punavirus*; s_*Escherichia* virus P1). This function is essential for regulating the *Caulobacter* cell cycle [38]. Interestingly, our analysis identified 7,255 contigs assigned to the *Caulobacter* genus, suggesting a potential interference with the biology of this taxon. Similarly, we found a gene encoding for a biofilm protein (BapA), also another protein known to interact with the lipopolysaccharide biosynthesis (*lpxH*) and a Nlpc-P60 domain (which has been related with host-microbe interactions [39–41], within sequences of the *Escherichia* virus P1. Additionally, we found a cellulose 1,4-beta-cellobiosidase within a *Tequatrovirus* contig and a chitinase (class I) within a contig assigned to *Bacillus* phage SPP1 (o_*Caudovirales*; f_*Siphoviridae*), both of which are enzymes associated with complex carbohydrate catabolism.

Despite detecting the aforementioned bacteriophages in our metagenomes, we hypothesize that the long-term environmental stability of the cave may have relaxed the selective pressure exerted by phage predation. In the absence of new viral threats, maintaining costly defense mechanisms may be less advantageous. To test this, we compared the abundance of phage-defense systems in the bins of the cave to those found in semblant environments from other locations (Supplementary Table 4, Supplementary Table 8), and we found that indeed the content on such defensive genes is lower in Movile (Wilcoxon rank-sum test, *p* = 5.3×10^-4^) (Figure 4b).

## Discussion

Our study provides a comprehensive evolutionary perspective on the Movile Cave microbiome, shedding light on how microbial communities persist and adapt under stable but resource-constrained conditions. By examining patterns of gene retention, pseudogenization, and HGT, we highlight the mechanisms that drive long-term genomic stability in an ecosystem isolated for ∼5.5 millions of years. In this context, the observed pseudogenization (loss of function) in the microbial communities of Movile Cave can be understood as part of a broader phenomenon of reductive genome evolution, which is suggested as a key adaptive mechanism [42–45]. While adaptive gene loss has been extensively documented in host-associated bacteria [46,47], studies on pseudogenes in environmental microbial communities remain scarce [48]. Pan-genome studies have demonstrated that even within species, the variation in gene content—often in the form of gene loss—can be widespread and plays a crucial role in adaptation [49,50]. Evidence suggests that environmental stressors like acidity, heat, or salinity might drive genome streamlining [51], but data from stable extreme environments are still limited.

The high prevalence of pseudogenization in genes related to translation and other housekeeping functions within the Movile Cave microbiome challenges classical models of genome evolution in free-living bacteria. While pseudogenization of niche-specific genes is a hallmark of genome reduction in obligate symbionts [47,52–53], the observed inactivation of core genes in a free-living community suggests that strong ecological constraints and long-term environmental stability may drive similar patterns in extreme ecosystems. One possible explanation is that the stability and nutrient limitation of the Movile Cave environment reduce the selective pressure on rapid growth and full translational efficiency. In such conditions, reduced metabolic and biosynthetic capacity may be advantageous, favoring genome streamlining even at the cost of partial loss of translational fidelity or repair mechanisms. The redundancy of essential functions across syntrophic microbial consortia may also buffer the impact of these losses, allowing for communal complementation of critical functions [14, 54]. Additionally, the pseudogenization of multiple housekeeping genes, typically used for phylogenomic reconstruction [37], highlights the need to reassess the assumptions of essentiality and evolutionary conservation in extremophile microbiomes. These findings provide novel insight into microbial adaptation in nutrient-poor, isolated ecosystems and underscore the potential for convergence between genome reduction processes in endosymbiotic and extremotrophic contexts.

The high microbial densities reported in Movile Cave (e.g.: up to 10⁷ CFU/g in microbial mats [55]) contrast with the expected evolutionary dynamics of small populations, where loss-of-function mutations are more likely to fix due to genetic drift [56]. In large populations, gain-of-function mutations typically dominate due to their higher fitness benefits and clonal interference preventing the fixation of most loss-of-function variants [56–58]. The prevalence of pseudogenes and the scarcity of singletons or horizontally acquired genes in Movile Cave suggests that other evolutionary pressures—such as long-term environmental stability and the energetic cost of maintaining superfluous genes— may override the classical population-size-driven dynamics of adaptive evolution. The lack of evidence for relaxation of purifying selection admits two complementary interpretations. First, genes that became superfluous in the cave environment have likely undergone pseudogenization and subsequent loss over evolutionary time, resulting in a core genome composed predominantly of genes whose functions remain essential under the prevailing ecological constraints. Second, for these retained core genes, the lack of clear signatures of relaxed purifying selection suggests that prolonged isolation has not been accompanied by a substantial reduction in effective population size.

In stable environments, where selection is expected to favor gene loss and the emergence of metabolic interdependencies, the persistence of high inter-bin redundancy challenges the predictions of the Black Queen Hypothesis [14]. This hypothesis posits that long-term selective pressures should promote a division of labor, with some taxa losing costly functions while relying on others to maintain them. One possible explanation is that, given the high microbial loads in the cave, competition may also serve as a counterbalancing force to functional specialization, favoring metabolic autonomy over reliance on communal functions. Our results support this idea: while adaptive gene loss has reduced functional potential at the individual genome level, community-wide redundancy remains high and may help maintain ecosystem stability in this unique environment. Notably, we observed that functions undergoing pseudogenization tend to be those with higher redundancy, suggesting that functions supported by multiple gene copies are more likely to tolerate gene degradation. In contrast, no pseudogenization events were detected among functions with extremely low redundancy (e.g., single-copy genes conserved across bins), highlighting selective constraints to preserve essential processes within the community. Pseudogenization may thus represent a fine-tuning mechanism to reduce excess redundancy in metabolically costly functions, without fully compromising the functional repertoire required for ecosystem persistence.

This cave harbors several yet undescribed microorganisms; however, the proportion of novel genetic material within their genomes was relatively low, as evidenced by the limited presence of singletons and HGT events. Similarly, a recent comparative study on “*Candidatus* Thiovulum stygium,” binned from DNA sourced in Movile Cave, revealed significant differences between this taxon and related “*Candidatus* Thiovulum” genomes, showing the lowest number of singletons in that pangenome analysis [59]. HGT has been suggested to facilitate bacterial adaptation to changing environmental conditions [60,61]. However, a recent study suggests that contrary to common assumptions, transitions to new ecosystems may actually lead to a decrease in HGT rates, as well as a strengthening of purifying selection [62]. This challenges the traditional view that environmental changes necessarily increase HGT rates due to the adaptive benefits of gene acquisition. The study found that environmental transitions, rather than promoting the influx of new genetic material, are often associated with a reduction in HGT within microbial communities [62]. The long-term environmental stability of the cave over 5.5 million years suggests that its microbial communities have reached a fitness peak within a highly specialized niche, where selective pressures remain constant. In such stable ecosystems, new gene acquisitions—whether through mutation or HGT—are less likely to be retained unless they confer a significant selective advantage. Unlike in an unoccupied niche, where novel functions can facilitate colonization and adaptation, in an already optimized and densely populated system, new genetic material may be neutral or even maladaptive, leading to its elimination through purifying selection or genetic drift [56,63,64]. Additionally, the absence of external microbial influx further limits opportunities for HGT, reinforcing the genetic conservatism observed in the cave microbiome.

Viral AMGs are known to affect functional diversification of microbial communities [65–69]. For instance, it has been shown that viruses affect organic matter cycling and the composition of the microbiota of microbial mats [70]. Furthermore, viruses have contributed to the regulation of essential biogeochemical cycles in both oceanic [71] and hydrothermal vent ecosystems [72], underscoring their ecological significance. In long-term stable ecosystems like the Movile Cave, viral-host interactions may play a subtle yet significant role in shaping microbial adaptation. Here, we reveal that the virome of this cave is predominantly composed of *Caudovirales*, carrying AMGs that could influence the microbial ecology and evolution of this unique habitat. For instance, one such AMG found in a viral sequence is the GcrA cell cycle regulator. Dimorphic cell cycles are rare among prokaryotes, with *Caulobacter crescentus* being one of the best-studied examples [73]. This bacterium transitions from a motile lifestyle (swarmer cell) to a surface-associated state (stalked cell), a process mediated by the GcrA regulator, which acts as a σ70 cofactor driving a cascade of gene expression [38]. Considering all this, we posit that the dissemination of this AMG through viral infections could influence the dynamics of the microbial mat within the cave. Similarly, we identified AMGs that affect the metabolism of molecules (e.g., BapA, lipopolysaccharides, cellulose) involved in the structure of biofilm matrixes [74,75], which may contribute to the maintenance of the microbial mat. In any case, considering that we found a slightly lower content of phage-defense systems in the Movile genomes, the effect of viruses in the dynamics of these microbiomes might be lower than in other environments due to the lack of exposure to novel phages or to the prolonged adaptation to them.

Finally, we propose the description of two novel taxa: “*Candidatus* Edaphobacter cavernae” and “*Candidatus* Sedimentibacter cavernae”. This is the first report of species from both genera thriving in a chemolithotrophic cave environment. All the described *Edaphobacter* species (*Acidobacteriota*) have been isolated from forest soils or lichen [76]. Interestingly, *Edaphobacter aggregans* Wbg-1^T^ was originally recovered from a co-culture with a slime-producing methanotrophic bacterium (*Methylocella silvestris*) established from calcareous forest [77,78]. In contrast, certain type strains from *Sedimentibacter* species have been isolated from methanogenic water sediments [79,80].

### Conclusions

The microbiome of this chemolithotrophic ecosystem that has been isolated from external influences for 5.5 Myr is largely specific. The long-term isolation has fostered a highly specialized microbial community, with evolutionary dynamics dominated by the selective retention of essential genes, the purging of non-essential functions, and a remarkably restricted horizontal gene flow. This restricted genetic exchange is likely driven by the lack of fluctuating selective pressures, the cave’s extreme ecological niche, and the absence of foreign microbiomes. Surprisingly, despite these pressures, we found no evidence of a significant loss of functional redundancy at the community level, challenging some predictions of evolutionary theory. However, at the genomic level, we detected signs of pseudogenization and erosion even among core housekeeping genes, suggesting that long-term ecological stability may allow the relaxation of purifying selection, even for traditionally conserved functions. This decoupling between individual genome erosion and community-level functional stability points to a possible buffering effect provided by the microbial consortium as a whole. Additionally, our findings highlight the impact of the virome on the exchange of ecologically relevant genes within the Movile Cave. These results offer an unprecedented view of microbial evolution in a stable, isolated environment, highlighting the adaptability of life under extreme conditions and the delicate balance between gene loss, functional redundancy, and ecological stability.

### Protologue

#### Description of “*Candidatus* Edaphobacter cavernae”

“*Candidatus* Edaphobacter cavernae” (ca.ver′nae. L. gen. n. *cavernae* of a cavern). It is proposed the genome sequence of dastool_concoct.19 (JAWQJF000000000) be considered as a reference sequence for ‘Ca. E. cavernae’ in the absence of biological resource material. The genome of the species has been recovered from a floating mat internal lake in the first limited oxygen airbell of a 5.5 Myr trapped cave.

This species contains genes for the assimilation of sulphite (EC 1.8.1.2), dehydrogenation of formate (EC 1.17.1.9) and bifunctional oligoribonuclease and PAP phosphatase (EC 3.1.3.7; EC 3.1.13.3). At the same time, this species contains genes for the utilization of the following carbon sources: cellobiose, ethanol, galactose, glucose, maltose, ribose, sucrose and trehalose.

The reference strain dastool_concoct.19 is associated with a cave environment (GenBank accession number: JAWQJF000000000); 16S rRNA gene accession number is OR689336. The genome size is around 4.7 Mbp and the G+C content is 57.2%.

#### Description of “*Candidatus* Sedimentibacter cavernae”

“*Candidatus* Sedimentibacter cavernae” (ca.ver′nae. L. gen. n. *cavernae* of a cavern). It is proposed the genome sequence of dastool_metabat2.39 (JAWQJG000000000) be considered as a reference sequence for ‘Ca. E. cavernae’ in the absence of biological resource material. The genome of the species has been recovered from a floating mat internal lake in the second limited oxygen airbell of a 5.5 Myr trapped cave.

This species contains genes for oxidoreductases (EC 1.2.7.1), and bifunctional oligoribonuclease and PAP phosphatase (EC 3.1.3.7; EC 3.1.13.3). At the same time, this species contains genes for the utilization of the following carbon sources: alanine and glutamate.

The reference strain dastool_metabat2.39 is associated with a cave environment (GenBank accession number: JAWQJG000000000). The genome size is around 2.4 Mbp and the G+C content is 32.4%.

## Methods

### Sample collection

The Movile Cave is located in Romania, close to the Black Sea (43.825487N; 28.560677E). This environment is completely driven by chemoautotrophy [32–35], resembling life in deep-see hydrothermal vents [28]. Its water and air temperature are about 20°C, with variable chemical composition depending on the site, with remarkably high levels of H_2_S, CH_4_ and NH_4_^+^ (see extensive details of the cave elsewhere [28]). Sampling was performed during one single visit in August 2014. In recognition of the need to protect this unique ecosystem, which relies heavily on the cave’s distinctive environmental conditions, stringent visiting protocols were implemented to prevent any potential external contamination. Samples of cave sediments and microbial mats, either floating on the water or collected from the cave walls, were carefully gathered into sterile plastic vials and tightly sealed to prevent gas exchange. These samples were obtained from airbells I and II as well as the lake room (Figure 1) and promptly transported to the Czech Republic, where they were frozen (-25°C) until further processed. Because the temperature in the sampling site was 18-21°C, we maintained this temperature during transport.

### DNA extraction and shotgun sequencing

Total DNA was extracted using 200 mg of each sample with the FastDNA™ SPIN Kit for Soil (MP Biomedical) according to the manufacturer’s protocol.

Libraries for metagenome sequencing were prepared with the NEBNext Ultra II DNA Library Prep Kit for Illumina (New England BioLabs), following the manufacturer’s instructions. Samples were pooled in equimolar volumes and sequenced in a HiSeq 2500 system (2 × 250 bp) (Illumina) at Brigham Young University Sequencing Centre, USA.

### Metagenomic sequences analyses

The raw reads were cleaned as described previously [40]. Briefly, we used Trimmomatic (v0.39) [81] for adapter removal. Then, we used FastX-Toolkit scripts (http://hannonlab.cshl.edu/fastx_toolkit/index.html) to trim with a Phred score < 30, and biopieces (http://www.biopieces.org) to remove sequences shorter than 50 bp. We normalized the reads by copy number with khmer [82]. We assembled the metagenomic contigs with Megahit (v1.2.9) [83]. When needed, we mapped the cleaned reads to the assembly with Bowtie2 [84] and converted the SAM files to BAM format with Samtools (v1.9) [85].

### Recovery of genomes and bins

We recovered metagenomic bins with 3 binning algorithms: MaxBin2 (v2.2.7) [86], MetaBAT2 (v2.15) [87], and Concoct (v1.1.0) [88]. Then, we de-replicated and generated consensus bins with DAS-Tool (v1.1.2) [89].

We performed a quality check and taxonomic assignment of the bins as previously done [19]. We estimated the contamination levels of the bins with CheckM (v1.1.3) [90], and the genome completeness with BUSCO (v5.4.3) [91]. Bins with <50% completeness and/or >10% contamination were discarded. To dereplicate the genomes, we used dRep (v3.4.0) [92] with a 95% ANI threshold (-sa 95) and a minimum completeness of 50%. All genomes were classified as distinct, and no genomes were clustered as redundant. We then assigned the taxonomy of bins with the GTDB-Tk program (v2.1.1) [93] (“classify_wf” command). This workflow uses the closest average nucleotide identity (ANI) value to locate the user strain into the closest species in the GTDB. Bins with completeness >90% and contamination <10% were termed metagenome-assembled genomes (MAGs).

### (Meta)genome functional analyses and comparative genomics

We annotated the functions of the genes/proteins with KofamscanScan (locally installed), KofamKOALA (webserver) [94] and Eggnog-Mapper (v2.1.9) [95]. We searched for antimicrobial- and stress-resistance genes with AMRFinderPlus [96]. We used GapMind to unravel the potential ability to catabolize small carbon sources of the predicted MAGs, considering only the nutrients with high or medium confidence according to the output [97].

We conducted comparative genomic analyses involving the 4 MAGs from the cave characterized at the genus or species level. As a reference for these analyses, we downloaded genomes and MAGs belonging to each of the 4 genera (*Sedimentibacter*, *Edaphobacter*, *Sulfurospirillum* and *Desulfuromonas*) from the IMG-JGI portal (https://img.jgi.doe.gov/), NCBI (https://www.ncbi.nlm.nih.gov/), and from the “Genomes from Earth’s Microbiomes (GEM) catalog” [98] (Supplementary Table 3). The comparative genomic analyses were done with Anvi’o [99], where genomes were first reformatted to standardize contig names and filtered to retain sequences longer than 200 bp. Within the Anvi’o scripts, gene prediction was performed with Prodigal [100] before generating contig databases, which were subsequently annotated using NCBI COGs and HMM searches. Pangenome analyses were conducted with default parameters and additional configurations for hierarchical clustering and minimum gene occurrence thresholds. Genomic similarity was assessed using pyANI [101]. We focused on the differential abundance of unique genes or singletons— those that remain unclustered in the pangenome analysis due to insufficient sequence similarity or alignment coverage thresholds during DIAMOND-based protein similarity searches and MCL clustering—as they likely represent lineage-specific adaptations or horizontally acquired functions that distinguish the cave microbiota from related taxa. Unlike core or accessory genes, which may be shared among multiple strains and contribute to conserved functions, singletons can provide insights into the selective pressures and ecological interactions shaping microbial life in the cave. Their presence may reflect niche-specific innovations, acquisition through horizontal gene transfer (HGT), or genomic drift due to prolonged isolation.

We used MetaCHIP (v1.10.10) [102] to search for horizontal gene transfer (HGT) events among genome bins recovered from the environmental metagenome. MetaCHIP predicts HGT by performing an all-against-all BLASTN search on predicted open reading frames (ORFs) from input genomes. It then compares the best matches across different groups of genomes, identifying putative HGT candidates based on higher identity scores from non-self groups. The candidates are further validated by analyzing their flanking regions and using a phylogenetic approach for corroboration. Based on the similarity between each pair of shared genes, we classified the HGT events as recent (>99% nucleotide similarity) or ancient (60-98% similarity) [102].

We compared the proportion of HGT events found in the cave and those likely occurred in similar microbiomes from other environments. We downloaded all the bins (>50% completeness, <10x contamination) recovered from 10 different metagenomes from hydrothermal vents, microbial mats, freshwater, sediments and groundwater (Supplementary Table 4) that are available at the GEM database [98]. In total, we used 712 external bins (33 – 218 bins/study) belonging to 408 taxa (23 – 110 taxa/study) as by the identification with GTDB-tk [93]. We then used MetaCHIP separately with each subset of bins from each microbiome. Beyond comparing the total HGT events identified, we relativized that count to the total number of taxa per study, since the transfer of genes was searched by comparing genera or higher taxonomic levels [102].

We measured the pseudogene content with Pseudofinder (v1.1.0) [103] (diamond search) over the genomes annotated with prokka (v1.14.6) [104] (‘--rfam --compliant’ flags), using as a reference the Swiss-Prot database retrieved from the NCBI (https://ftp.ncbi.nlm.nih.gov/blast/db/FASTA/swissprot.gz; accessed on November-2023). Default settings were used for all other parameters. Then, the level of genome pseudogenization was compared among groups of genomes by relativizing the total count of pseudogenes by the count of CDSs per genome. We also analyzed the total count of pseudogenes evidenced from significant matches to database sequences outside gene annotations, focusing on intergenic pseudogenes as they are more likely to represent degenerating gene sequences compared to other pseudogene classes [48]. Pseudofinder is configured to omit contig termini from the BlastX analysis of intergenic sequences, minimizing false-positive pseudogene predictions near assembly boundaries. The GFF file generated by Pseudofinder contained the classification into intergenic or other different pseudogenes. The same workflow was applied to all the genomes to avoid biases.

As an indicative of defense against bacteriophages, we searched for genes related with the defense against these viruses with DefenseFinder (v1.2.0) [105]. This tool allows to perform quick searches against HMM models of known anti-phage systems. The viral contigs were taxonomically identified with MMSeqs2-taxonomy (v13.45111) [106] against the UniProt database (UniProtKB/Swiss-Prot Release 2021_03) [107].

### Selection analyses

For the analysis of selection patterns in microbial genomes, we first used PPanGGOLiN (v1.2.105) [108] in order get the core genes for each group of tested genomes. The parameters for sequence similarity and coverage were set to 80%. The resulting pangenome was used for subsequent analysis of the core genome, which was partitioned for DNA sequences and aligned using the PPanGGOLiN msa command, which relies on mafft (v7.520) [109] with default options to perform the alignment. The generated multiple sequence alignment (MSA) was used for phylogenetic tree construction and selection analyses. To ensure the integrity of the sequence data for downstream analyses, we performed a series of data preprocessing steps. The sequences were reformatted into a standardized FASTA format using additional custom scripts. Stop codons were removed from the alignments using a HyPhy ‘cln’ command (v2.5.62) [110]. Sequence names were further cleaned to eliminate any invalid characters, such as periods, which could interfere with processing.

Once the alignments were cleaned, we used HyPhy (v2.5.62) [110] to perform selection analysis using two distinct methods: aBSREL and RELAX. The aBSREL method was employed to assess positive selection on individual branches of the phylogenetic tree, and the RELAX method was used to evaluate whether purifying selection was relaxed or intensified along specific branches. HyPhy’s ability to analyze selection at specific codon positions within genes—rather than analyzing the entire gene—provides a finer resolution for detecting variation in selective pressures across different sites. Both analyses were applied with the appropriate phylogenetic tree, which was constructed using RAxML (v8.2.12) [111] based on the alignment obtained with PPanGGOLiN, using a concatenation of all core genes (via the –phylo command in the msa step). The tree was built with the General Time Reversible model with Gamma-distributed rate heterogeneity (GTRCAT) and a random seed was set using the -p 123 flag to ensure reproducibility of the results. The phylogenetic tree was rooted, and selection tests on branches were conducted by labeling the test genome as “Foreground” and the sister genomes within the same branch as “Background” in the tree. Results from both selection analyses (aBSREL and RELAX) were processed using custom Python scripts to extract and classify the results. The processed results were used to identify potential genes under adaptive selection in the microbial communities.

### Statistical analyses

All statistical analyses were conducted in R. To assess the normality of the data, the Shapiro-Wilk normality test was performed using the shapiro.test function. Data that did not follow a normal distribution were analyzed using non-parametric methods. The Wilcoxon rank-sum test, applied through the wilcox.test function, was used to evaluate the differences in the number of pseudogenes and defense genes between groups. This test was chosen as it is suitable for comparing two independent groups when the data are not normally distributed.

### Phylogenies and Overall Genome Relatedness Indices (OGRIs) measurements

We investigated the taxonomy of the MAGs following a workflow that encompasses phylogenomics and the calculation of the Overall genome relatedness indexes (OGRI) [112,113]. We built the phylogenomic trees with the UBCG (v3) tool (default settings) [114] which created alignments (with MAFFT), and trees (with RAxML) based on 92 housekeeping genes. Then, we visualized and customized the consensus tree in The Interactive Tree Of Life (iTOL) tool (v5) [115]. Digital DNA–DNA hybridization (dDDH) values were calculated *in silico* with the Genome-to-Genome Distance Calculator (GGDC 2.1) using the blast method (https://tygs.dsmz.de/) and the recommended formula #2, which is independent of genome length and appropriate for incomplete genomes [116]. Average Nucleotide Identity values based on blast comparisons (ANIb) were calculated with PYANI (v0.2.10) [117]. We used both genome similarity indexes (dDDH and ANIb) to confirm species taxonomic assignments. Values of 70% dDDH and 95-96% ANIb are considered as thresholds for species delimitation [118,119]. Individual analysis of whole-genome phylogeny of MAGs was generated at Type (Strain) Genome Server (TYGS) [120].

## Declarations

### Authors contributions

Zaki Saati-Santamaría (Conceptualization, Formal analysis, Investigation, Software, Methodology, Visualization, Writing – original draft, Writing – review and editing); Alena Nováková (Investigation, Methodology); Lorena Carro (Investigation, Methodology); Fernando González-Candelas (Conceptualization, Methodology, Writing – review and editing); Jaime Iranzo (Conceptualization, Methodology, Writing – review and editing); Alexandra Maria Hillebrand-Voiculescu (Investigation, Resources); Daniel Morais (Formal analysis, Methodology); Tomáš Větrovský (Formal analysis, Methodology); Petr Baldrian (Resources, Funding acquisition, Writing – review and editing); Miroslav Kolarik (Conceptualization, Funding acquisition, Project administration, Resources, Supervision, Writing – review and editing); Paula García-Fraile (Conceptualization, Methodology, Investigation, Supervision, Writing – review and editing).

## Acknowledgements

ZSS, LC and PGF acknowledge the funds received by the Regional Government of Castilla y León, Escalera de Excelencia CLU-2018-04, co-funded by the P.O. FEDER of Castilla y León 2014–2020. ZSS acknowledge a grant funded by Program EU Horizon Europe (HORIZON-TMA-MSCA-PF-EF; Grant n° 101090267), and a grant cofinanced by the European NextGenerationEU, Spanish “Plan de Recuperación, Transformación y Resiliencia,” Spanish Ministry of Universities, and the University of Salamanca (“Ayudas para la recualificación del sistema universitario español 2021-2022”). FGC was funded by projects PID2021-127010OB-I00 from Spanish Ministry of Science (MICIN) and CIPROM-2021-053 from Generalitat Valenciana. JI is supported by the Agencia Estatal de Investigación of Spain (Grant No. CNS2023-145430). TV was supported by the Czech Science Foundation (25-15251S), PB was supported by the Ministry of Education, Youth and Sports of the Czech Republic (MŠMT CZ.02.01.01/00/22_008/0004597). The sampling campaign occurred in the frame of the project “Cave invertebrates and microorganisms – their interactions, ecophysiological and morpho-functional adaptations” – an interacademic exchange between the Czech Academy of Sciences and the Romanian Academy.

## Data availability

All raw sequences obtained through metagenomic sequencing are available at the NCBI Sequence Read Archive (SRA) database under the Bioproject PRJNA1028180. The code of the sample filenames match to the names used in Supplementary Table 1. The genomic sequences (MAG) of the proposed *Candidatus* taxa are accessible in GenBank via the following accession numbers: JAWQJF000000000, JAWQJG000000000. All the genomes binned within this study (>50 completeness; <10% contamination; Supplementary Table 1) together with their proteomes are available at Zenodo (https://doi.org/10.5281/zenodo.10012125).

## Supplementary Data

Supplementary Data. File with a full description of the taxonomic and phylogenomic analyses performed over the metagenome-assembled-genomes (MAGs) resolved from Movile Cave.

Supplementary Table 1. Features of the assembled bins with >50% completeness level and <10% contamination.

Supplementary Table 2. Relative abundance of the metagenome-assembled-genomes (MAGs) from Movile Cave. The highest abundance of each MAG among all the samples is highlighted in bold.

Supplementary Table 3. Metadata for genomes and MAGs used in the comparative genomic analyses (Figure 3, Supplementary Figure 1, Supplementary Figure 2, Supplementary Figure 3).

Supplementary Table 4. Metadata of the bins and their original metagenomes used as external references to compare genome evolution events in Movile Cave bins. These bins were obtained from Nayfach et al. (2021) (https://portal.nersc.gov/GEM/). Different locations are distinguished by blue and white backgrounds.

Supplementary Table 5. Frequency of intact and pseudogenized occurrences of core and non-core functions in Movile Cave MAGs. The COG functions were retrieved from EggNOG-mapper annotations. Here we represent only the count of intergenic pseudogenes. As these are not predicted CDSs by genecalling algrithms, these cannot be directly annotated, so we used the COG annotation of the closest Swiss-Prot homologs employed as references in the Pseudofinder annotation. To count the number of COG functions that remain intact we annotated the 111 Movile Cave bins.

Supplementary Table 6. List of genes likely transferred horizontally from one bin to another. The table includes the nucleotide identity of each pair of homologous genes, as well as the annotation of each encoded protein, which in most cases is the same in both genes of each pair. The ‘Direction’ column indicates the putative donor bin (left) and recipient bin (right).

Supplementary Table 7. Horizontal gene transfer events detected among the bins from our study and those from microbiomes found in other environments. These bins were obtained from Nayfach et al. (2021) (https://portal.nersc.gov/GEM/). Further details are provided in Supplementary Table 4.

Supplementary Table 8: Defense-related genes identified in prokaryotic replicons using Defense-Finder. This table lists the detected defense genes, including their replicon of origin, unique identifiers, assigned gene names, and HMMER-based scoring metrics. Hit_status = the status of the component in the assigned system’s definition. See Supplementary Table 4 for details of each bin.

Supplementary Figure 1. Quality assessment of the recovered bins. This graph displays the quality of the metagenome-assembled genomes (MAGs) based on contamination (%) (X-axis), completeness (%) (Y-axis), and genome length (dot size). MAGs are shown in purple.

Supplementary Figure 2. Comparative genomic analysis of the *Desulfuromonas* sp. metagenome-assembled genome (MAG) from Movile Cave (dastool_maxbin2_145) alongside genomes and MAGs from other environments within the same genus (see Supplementary Table 3). Each circle represents the presence (dark color) or absence (light color) of gene clusters within each genome. The purple circle highlights the Movile Cave MAG. Shared ANIb values across all genomes are depicted in the purple heatmap. Left panel: Functional categories of the singletons found in the *Desulfuromonas* sp. MAG from this study, illustrating the metabolic traits associated with its ecological specialization in this environment. Notably, this genome contains the fewest singletons among all the analyzed genomes of this genus.

Supplementary Figure 3. Comparative genomic analysis of the *Sedimentibacter* sp. metagenome-assembled genome (MAG) from Movile Cave (dastool_metabat2.39) alongside genomes and MAGs from other environments within the same genus (see Supplementary Table 3). Each circle represents the presence (dark color) or absence (light color) of gene clusters within each genome. The purple circle highlights the Movile Cave MAG. Shared ANIb values across all genomes are depicted in the purple heatmap. Left panel: Functional categories of the singletons found in the *Sedimentibacter* sp. MAG from this study, illustrating the metabolic traits associated with its ecological specialization in this environment. Notably, this genome contains the fewest singletons among all the analyzed genomes of this genus.

Supplementary Figure 4. Comparative genomic analysis of the *Edaphobacter* sp. metagenome-assembled genome (MAG) from Movile Cave (dastool_concoct.19) alongside genomes and MAGs from other environments within the same genus (see Supplementary Table 3). Each circle represents the presence (dark color) or absence (light color) of gene clusters within each genome. The purple circle highlights the Movile Cave MAG. Shared ANIb values across all genomes are depicted in the purple heatmap. Left panel: Functional categories of the singletons found in the *Edaphobacter* sp. MAG from this study, illustrating the metabolic traits associated with its ecological specialization in this environment.

Supplementary Figure 5. Ratio of pseudogenes normalized by the total number of coding sequences (CDSs) per genome in both the bins obtained from Movile Cave metagenomes and those from other comparable environments.

Supplementary Figure 6. Redundancy of COG functions in the Movile Cave pseudogenes. The boxplots compare the copy number (i.e., redundancy) of COGs that include at least one pseudogenized gene versus those that do not include any predicted pseudogenes, and are split by COG categories. Since pseudogenes cannot be directly annotated, we used the COG annotation of the closest Swiss-Prot homologs employed as references in the Pseudofinder annotation.

Supplementary Figure 7. Total number of horizontal gene transfer (HGT) events detected among the bins from Movile Cave and those from microbiomes of other related environments (see Supplementary Table 4).

