## Supplementary Data for "Long-term evolution of prokaryotic genomes in a chemolithotrophic cave over 5.5 million years of isolation"

[1] Institute of Microbiology of the Czech Academy of Sciences, Vídeňská, Prague, Czech Republic
[2] Departamento de Microbiología y Genética, Universidad de Salamanca, Salamanca, Spain
[3] Institute for Agribiotechnology Research (CIALE), Villamayor, Salamanca, Spain
[4] Joint Research Unit "Infection and Public Health" FISABIO-University of Valencia, Institute for Integrative Systems Biology I2SysBio, Valencia, Spain
[5] Centro de Astrobiología (CAB), CSIC-INTA, Ctra. de Ajalvir Km 4, 28850 Torrejón de Ardoz, Madrid, Spain.
[6] Biospeology and Edaphobiology Department, *Emil Racoviţă* Institute of Speleology, Romanian Academy, str. Frumosă, no.31, sect.1, Bucharest 010986, Romania
[7]Group for Underwater and Speleological Exploration, str. Frumoasă, no. 31, Bucharest 010986, Romania
[8] Norwegian College of Fishery Science, UiT the Arctic University of Norway, Muninbakken 21, Tromsø 9019, Norway
[9] Biological Institute of São Paulo – Vila Mariana, Av. Conselheiro Rodrigues Alves 1252, São Paulo, SP 04014-002, Brazil
[10] Associated Unit USAL-CSIC (IRNASA), Salamanca, Spain
[†] Joint Senior Authors

---

### Supplementary Results:

#### Taxonomic identity of MAGs

Only a partial sequence of the 16S rRNA gene was recovered for one of the MAGs, concoct_19, revealing that its closest related species is *Edaphobacter dinghuensis* CGMCC 1.12997T (99.73% similarity). Its *gyrB* gene sequence (2,646 bp) shows the highest similarity (96–97%) to members of the genus *Edaphobacter*. This strain shares a dDDH value of 29.4% with *E. dinghuensis* CGMCC 1.12997T, confirming that concoct_19 represents a novel species, which is consistent with the phylogenomic analysis (Supplementary Figure). Based on these findings, we propose the name *Candidatus Edaphobacter cavernae*.

Genomic analysis of potential carbon source utilization suggests that this novel species can metabolize several compounds with high confidence, according to GapMind classification criteria. These include cellulose (bgl, MFS-glucose, flk genes), ethanol (ethanol dehydrogenase, *adh*, *acs* genes), galactose (Na+-galactose transporter, *galK*, *galT*, *galE*, *pgmA* genes), glucose (MFS-glucose, *glk* genes), maltose (*susB*, MFS-glucose, *glk* genes), ribose (BT2809, *rbsK* genes), sucrose (*ams*, MFS-glucose, *glk* genes), and trehalose (*treF*, MFS-glucose, *glk* genes). Additionally, the strain may utilize alanine, fructose, fucose, glycerol, mannose, proline, propionate, rhamnose, and threonine with moderate confidence

For metabat2_39, whole-genome phylogenetic analysis indicates that its closest related genera are quite distant (Supplementary Figure). The nearest species, *Sedimentibacter saalensis* DSM 13558$^T$, shares a dDDH value of only 21.5%. Furthermore, the dDDH values between metabat2_39 and other members of *Sedimentibacter* are even lower, supporting its classification as a novel species within this genus. We propose the name *Candidatus Sedimentibacter cavernae*. Carbon source utilization analysis suggests that this strain can metabolize alanine (*alsT* gene) and glutamate (*gltS*, *gdhA* genes) with high confidence, while asparagine, citrate, 2-oxoglutarate, and ribose are utilized with moderate confidence. These findings are consistent with previous observations for other *Sedimentibacter* species (Breitenstein et al., 2002).

One of the MAGs, maxbin2.145, lacks both the 16S rRNA and *gyrB* genes within its binned contigs. However, whole-genome phylogenetic analysis (Supplementary Figure) indicates that its closest relatives belong to the genus *Desulfuromonas*, with *Desulfuromonas acetexigens* DSM 1397$^T$ being the most similar strain (dDDH = 29.7%). This suggests that maxbin2.145 represents a novel species. However, due to the absence of genome sequences for other members of the genus, including the type strains of *Desulfuromonas chloroethenica*, *Desulfuromonas michiganensis*, and *Desulfuromonas carbonis*, its formal classification remains uncertain. Functional analysis reveals that this strain primarily utilizes compounds from the glyoxylate cycle, including acetate (*satP*, *ackA*, *pta* genes), fumarate (*dctM*, *dctP*, *dctQ* genes), L-malate (*dctM*, *dctP*, *dctQ* genes), and succinate (*dctQ*, *dctM*, *dctP* genes). Additionally, genes for the metabolism of ethanol, L-lactate, and serine were identified with moderate confidence.

For metabat2_12, whole-genome phylogenetic analysis suggests that its closest relatives include the type strain of *Rummeliibacillus suwonensis* (Supplementary Figure), with a dDDH value of 61.4%. However, this strain also shares the same dDDH similarity (61.4%) with other members of the genus and with *Mongoliibacter ruber*, the only species in the genus *Mongoliibacter* (Supplementary Figure). This ambiguity prevents a definitive classification until additional genome sequences from these genera become available. Carbon source utilization analysis indicates high confidence for alanine (*alsT* gene) metabolism, while acetate, aspartate, fumarate, and succinate are utilized with moderate confidence, aligning more closely with metabolic profiles observed in *Rummeliibacillus* species (Her & Kim, 2013).

Finally, for metabat2_89, phylogenomic analysis suggests that it is equally distant from genera belonging to different families within the phylum Actinomycetota, indicating that it likely represents a novel genus or family (Supplementary Figure). However, this observation may also be attributed to its relatively small genome size (2.4 Mbp) compared to closely related members of the phylum (5–10 Mbp), suggesting that the genome has not been fully recovered.

### References:

Breitenstein A, Wiegel J, Haertig C, Weiss N, Andreesen JR, Lechner U. Reclassification of *Clostridium hydroxybenzoicum* as *Sedimentibacter hydroxybenzoicus* gen. nov., comb.

nov., and description of *Sedimentibacter saalensis* sp. nov. *Int J Syst Evol Microbiol*. 2002 May;52(Pt 3):801-807. doi: 10.1099/00207713-52-3-801.

Her, J., & Kim, J. (2013). Rummeliibacillus suwonensis sp. nov., isolated from soil collected in a mountain area of South Korea. *Journal of Microbiology*, *51*, 268-272.

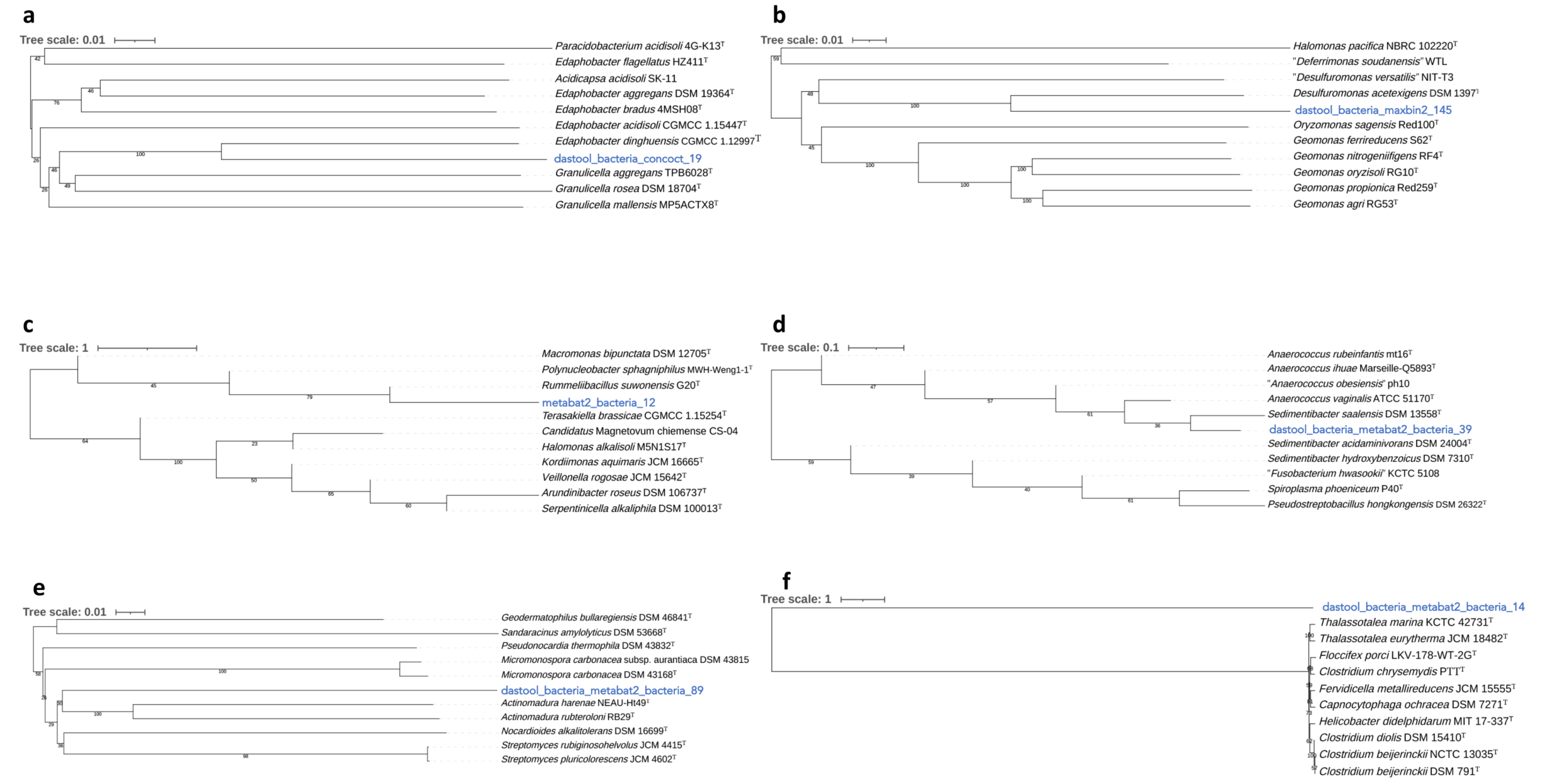


**Supplementary Figure**. Whole genome phylogenetic trees generated with TYGS of the HG-MAGs of the study (**a**) dastool_bacteria_concoct_19; **b**) dastool_bacteria_maxbin2_145; **c**) metabat2_bacteria_12; **d**) dastool_bacteria_metabat2_bacteria_39; **e**) dastool_bacteria_metabat2_bacteria_89; **f**) dastool_bacteria_metabat2_bacteria_14) and the closely related type strains with available genomes. Each tree was inferred with FastME from GBDP distances calculated from the genome sequences. Branch lengths are scaled in terms of the GBDP distance formula d5 and the numbers above the branches indicate the GBDP *pseudo*-bootstrap support values from 100 replications.
