## Supplementary figures and images for "Long-term evolution of prokaryotic genomes in a chemolithotrophic cave over 5.5 million years of isolation"

### Supplementary Figure 1

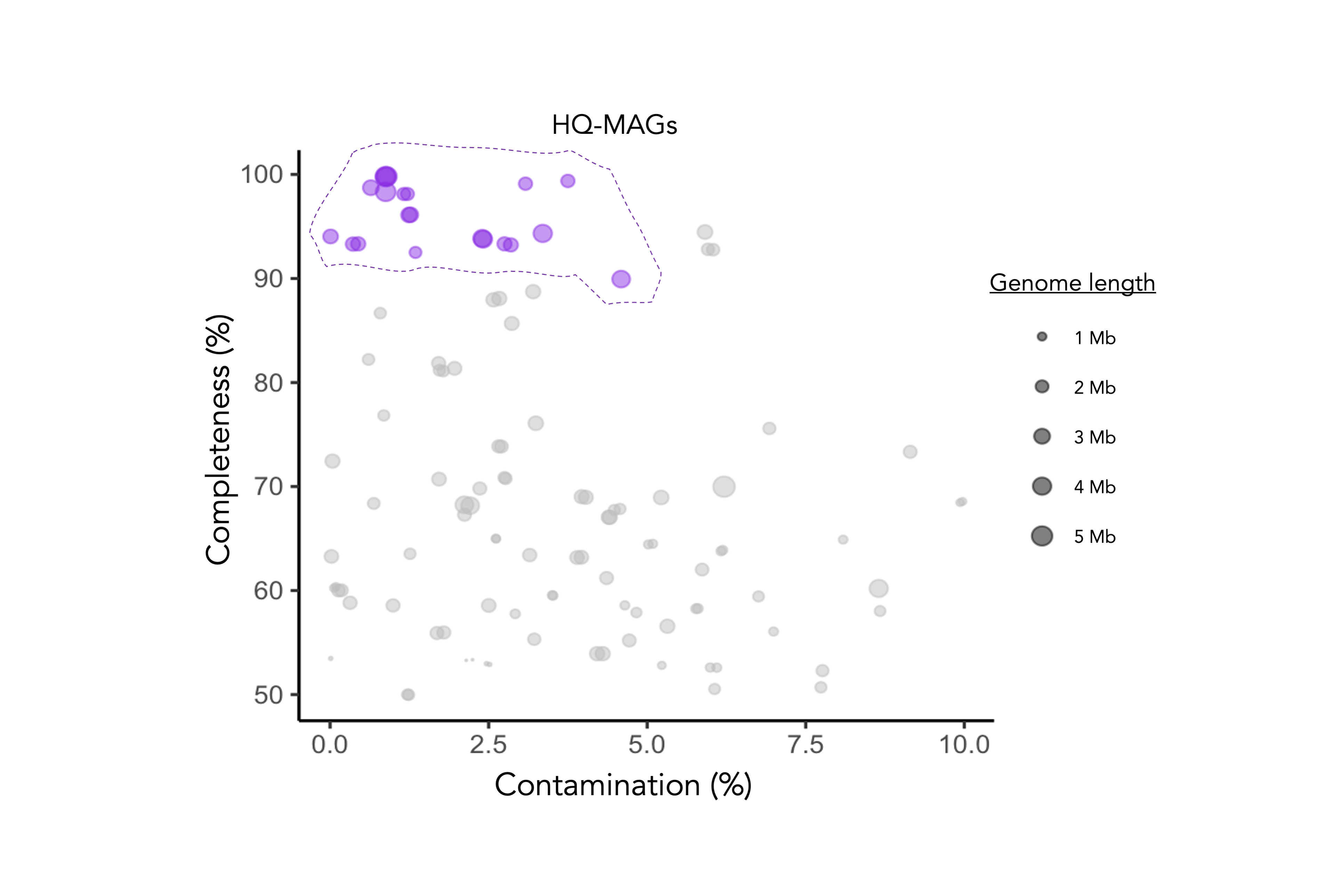

### Supplementary Figure 2

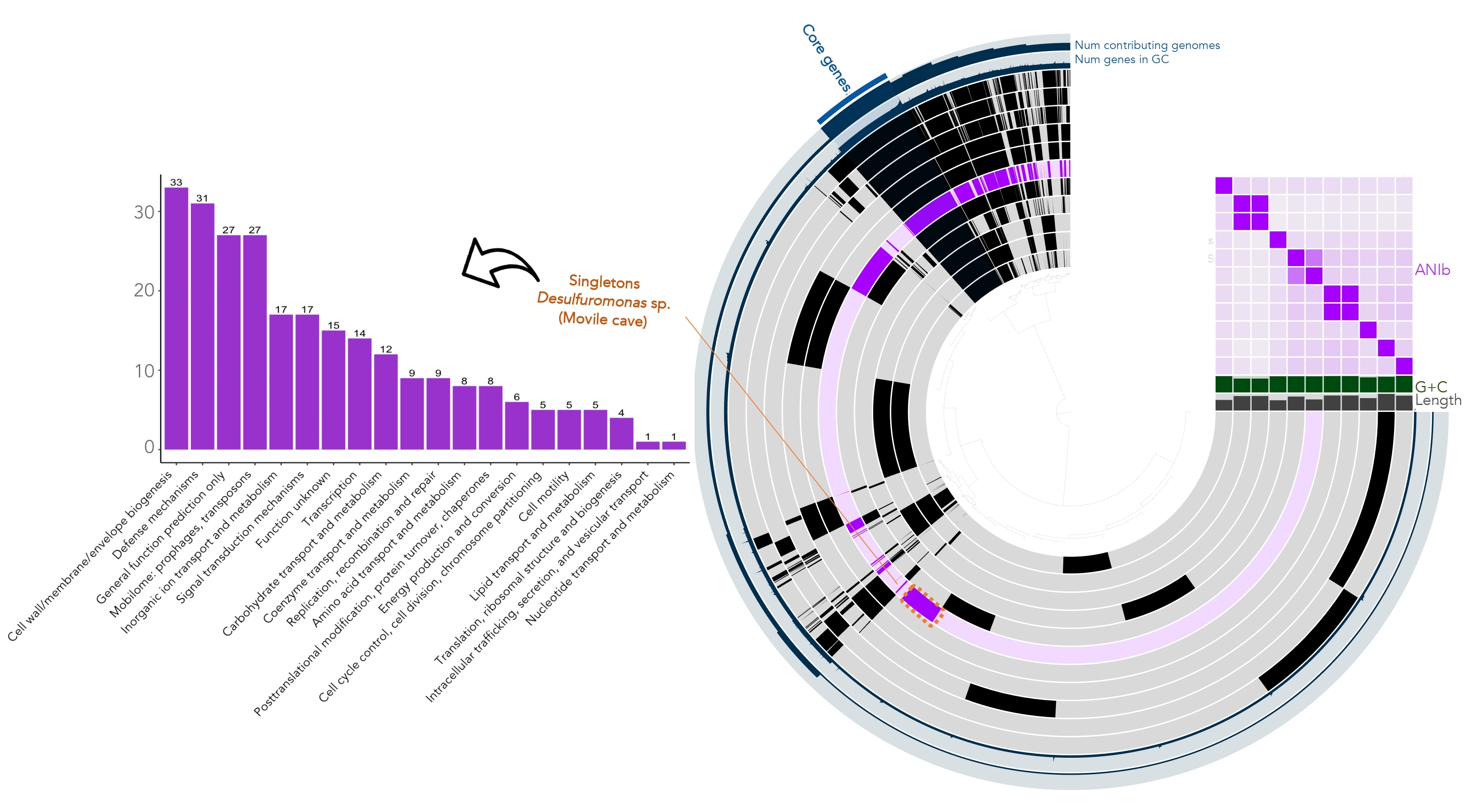

### Supplementary Figure 3

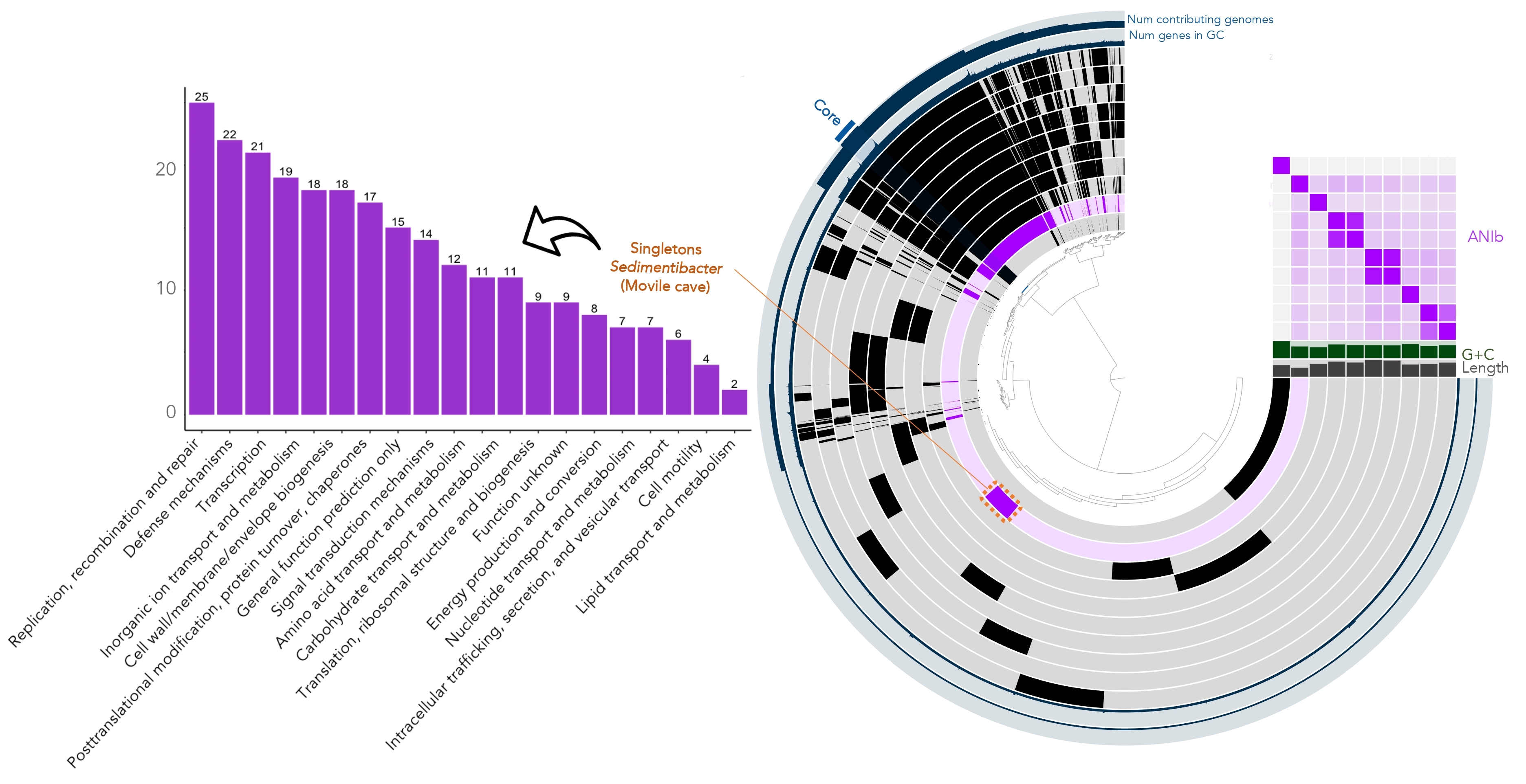

### Supplementary Figure 4

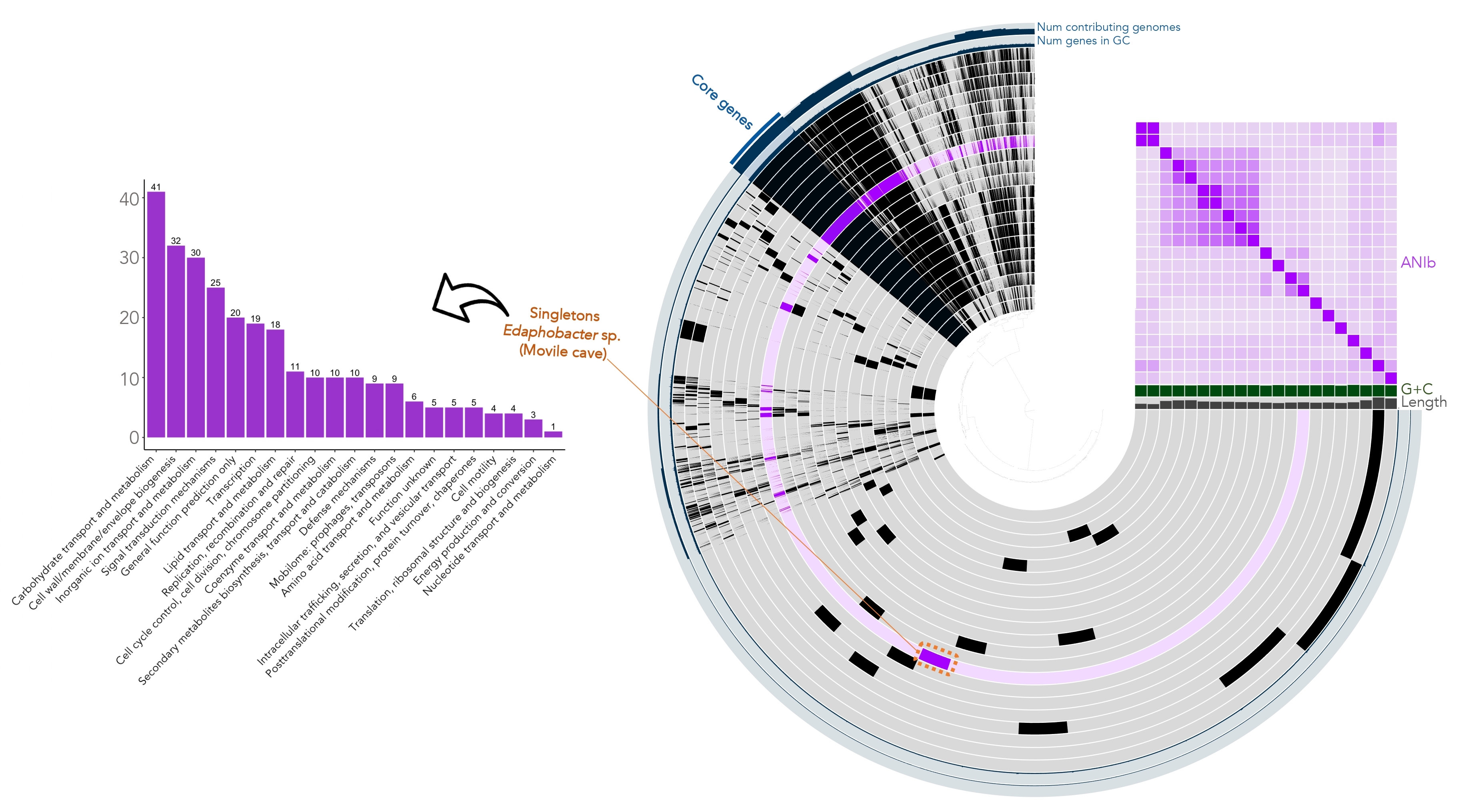

### Supplementary Figure 5

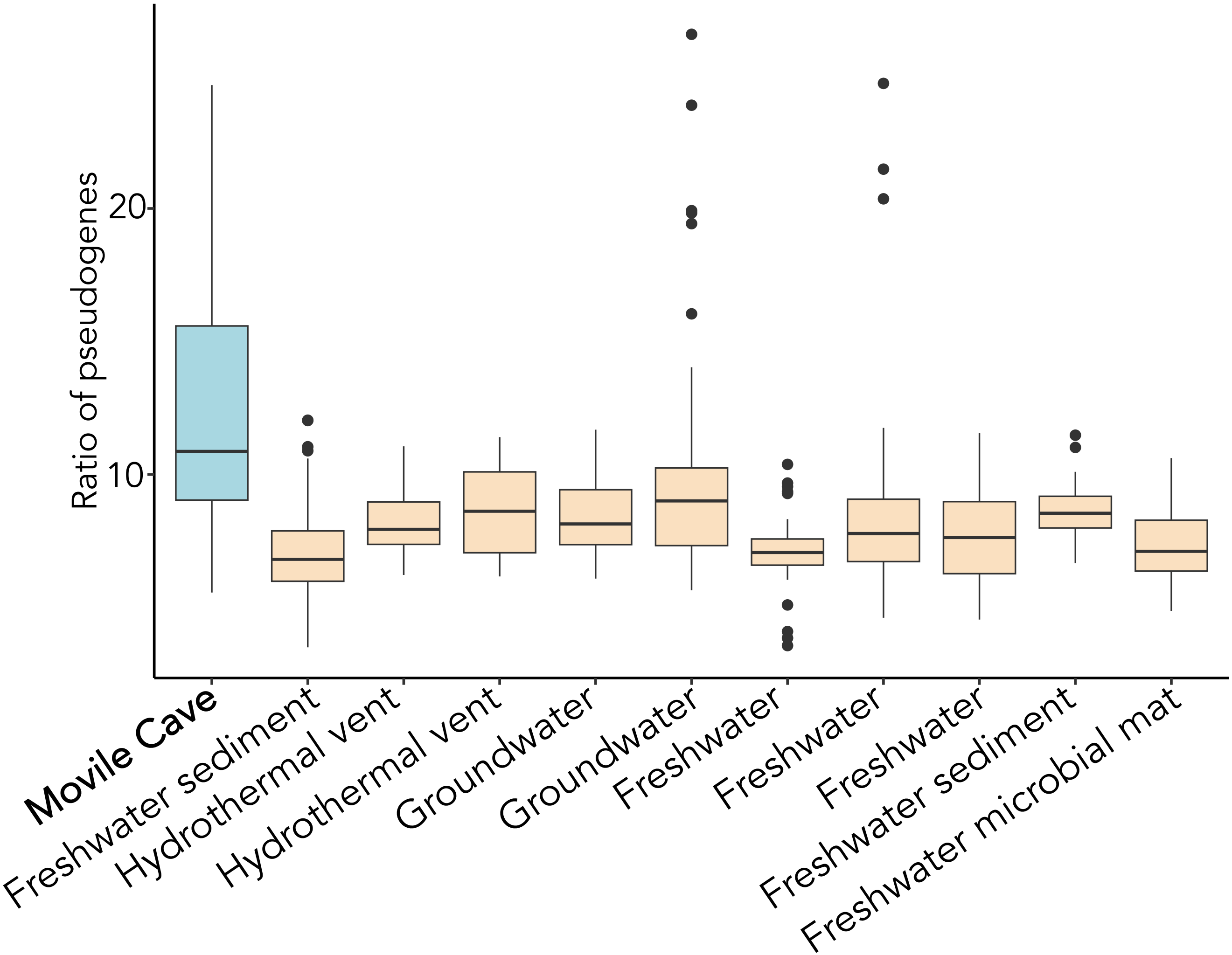

### Supplementary Figure 6

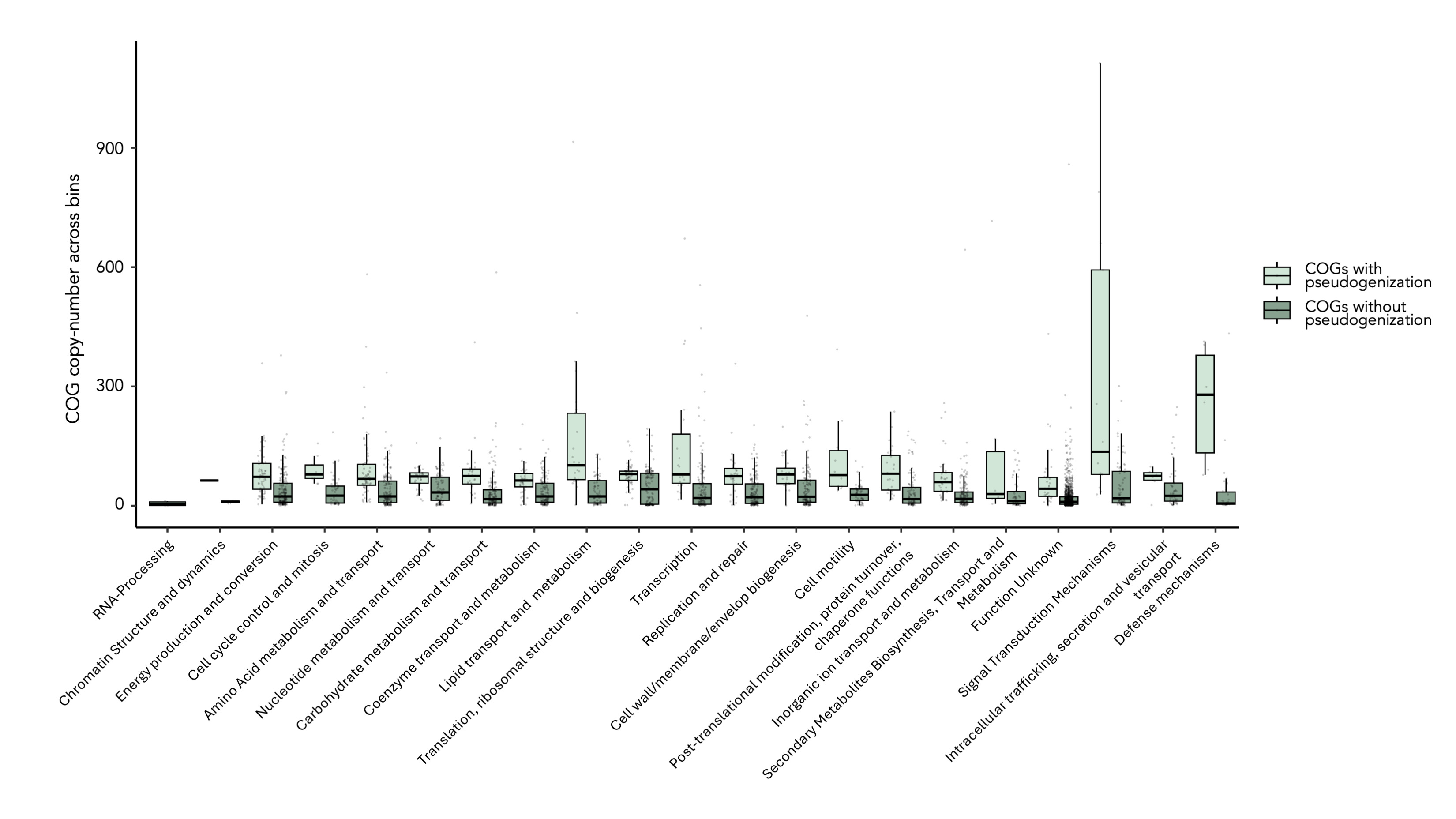

### Supplementary Figure 7

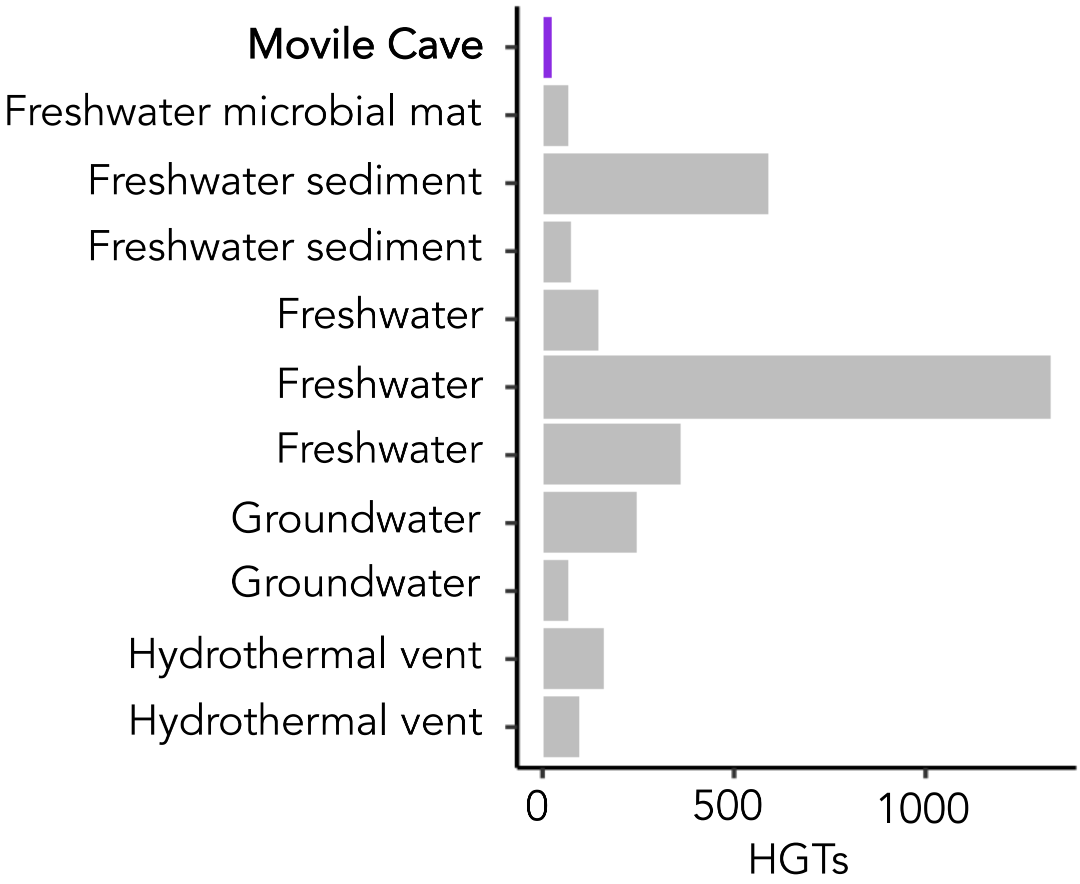
